# A valid protective immune response elicited in rhesus macaques by an inactivated vaccine is capable of defending against SARS-CoV-2 infection

**DOI:** 10.1101/2020.08.04.235747

**Authors:** Hongbo Chen, Zhongping Xie, Runxiang Long, Shengtao Fan, Heng Li, Zhanlong He, Kanwei Xu, Yun Liao, Lichun Wang, Ying Zhang, Xueqi Li, Xingqi Dong, Tangwei Mou, Xiaofang Zhou, Yaoyun Yang, Lei Guo, Jianbo Yang, Huiwen Zheng, Xingli Xu, Jing Li, Yan Liang, Dandan Li, Zhimei Zhao, Chao Hong, Heng Zhao, Guorun Jiang, Yanchun Che, Fengmei Yang, Yunguang Hu, Xi Wang, Jing Pu, Kaili Ma, Lin Wang, Chen Chen, Weiguo Duan, Dong Shen, Hongling Zhao, Ruiju Jiang, Xinqiang Deng, Yan Li, Hailian Zhu, Jian Zhou, Li Yu, Mingjue Xu, Huijuan Yang, Li Yi, Zhenxin Zhou, Jiafang Yang, Nan Duan, Huan Yang, Wangli Zhao, Wei Yang, Changgui Li, Longding Liu, Qihan Li

## Abstract

With the relatively serious global epidemic outbreak of SARS-CoV-2 infection, public concerns focus on not only clinical therapeutic measures and public quarantine for this disease but also the development of vaccines. The technical design of our SARS-CoV-2 inactivated vaccine provides a viral antigen that enables the exposure of more than one structural protein based upon the antibody composition of COVID-19 patients’ convalescent serum. This design led to valid immunity with increasing neutralizing antibody titers and a CTL response detected post-immunization of this vaccine by two injections in rhesus macaques. Further, this elicited immunoprotection in macaques enables not only to restrain completely viral replication in tissues of immunized animals, compared to the adjuvant control and those immunized by an RBD peptide vaccine, but also to significantly alleviate inflammatory lesion in lung tissues in histo-pathologic detection, compared to the adjuvant control with developed interstitial pneumonia. The data obtained from these macaques immunized with the inactivated vaccine or RBD peptide vaccine suggest that immunity with a clinically protective effect against SARS-CoV-2 infection should include not only specific neutralizing antibodies but also specific CTL responses against at least the S and N antigens.

## Introduction

Since the end of last year, a new species of coronavirus, a contagious agent capable of causing acute and severe respiratory infection and pneumonia through airborne transmission, has rapidly caused global public concern and even popular panic (1–3); the specific coronavirus was named SARS-CoV-2 by the World Health Organization (WHO) (4). Updated data have reported more than 3 million infection cases in the adult age group (5), with a death rate of approximately 2-10%, mostly in the elderly population (6, 7). With its high pathogenicity and infectivity and its spread in 200 countries and areas in a short time (8–10), not only multiple measures for clinical treatment and cure and public isolation but also vaccine development are urgently needed, and vaccine development could play a proactive role in controlling this epidemic (11). However, as SARS-CoV-2 is a new viral agent with an unknown infection mechanism and an unclear interaction with the immune system, vaccine development should first answer some basic questions, including characterization of antigenic component of this virus and its immunogenicity; validation of the protective immunity elicited by the viral antigen via certain immune processes; and whether the process of eliciting immunity to a viral antigen may be associated with immunopathogenesis during viral infection, similar to many concerns related to antibody-dependent enhancement (ADE) (12, 13). Based on these considerations, the work here raised the following inferential idea: the fact that SARS-CoV-2 infects cells through its spike (S) protein undergoing membrane binding with angiotensin converting enzyme 2 (ACE2) molecules in the cell membrane indicates the S protein is a major viral antigen that elicits a neutralizing antibody response (14, 15), while the nucleocapsid (N) protein, the other major structural component, acts as an antigenic stimulator of the innate immune response through its recognition as a pathogen-associated molecule pattern (PAMP) by cellular pattern recognition receptors (PRRs) in epithelial cells (16, 17). If this is the case, the N antigen should be significant in the study of viral vaccines, and the antibodies against the N protein may play a role in the antiviral immunity expected in vaccine development. Logically, these N-specific antibodies should be considered with the S-specific antibodies to be related to ADEs, which might exist in SARS-CoV-2-infected individuals (12). Our work describes a SARS-CoV-2 inactivated vaccine developed based on the above deduction and suggests a characterized immune response capable of defending against viral attack in rhesus macaques immunized with this vaccine, while the possible ADEs induced by existing antibodies against the S and N proteins are negated, as indicated by pathological observation and immunological detection. Conclusively, the inactivated vaccine is identified as having application potential.

## Results

### The antigenicity of the SARS-CoV-2 inactivated vaccine interacting with convalescent serum from COVID-19 patients

Previous studies of viral inactivated vaccines have indicated that the capacity of vaccines to elicit valid antiviral immunity in immunized individuals depends upon the viral antigenicity that is required to have viral antigenic components and structures displayed to the immune system of the body during the natural infectious process (18–20). With little knowledge about SARS-CoV-2, our inactivated vaccine created using a specific inactivating process was developed and investigated for its antigenicity by studying its interaction with convalescent serum, which was inferred to reflect, to some extent, the valid immune reactions of recovered individuals against viral infection. The results of a series of experiments were helpful in developing this vaccine. First, dozens of convalescent serum samples showed a neutralizing effect on the seed virus of the vaccine with varied titers of 1:16-256 (Fig. 1a), which suggests a potential quantized responsiveness of immunity leading to recovery in infected individuals. Furthermore, 2D electrophoresis and immune blotting using the convalescent serum suggested that not only the S protein but also the N protein and other proteins in this inactivated vaccine were recognized by the serum, which was more extensive recognition than that observed by blotting using a mAb against the S or N protein (Fig. 1b), and importantly, the anti-N antibody appeared to be an ascendant component. This result suggests that convalescent serum contains more antiviral antibodies than simply neutralizing antibodies and that these additional antibodies may play multiple functions in antiviral immunity. Our work using ELISA plates coated with the purified S or N protein or whole virions of the inactivated vaccine detected the convalescent serum and showed an interesting relationship between the anti-S, anti-N and anti-whole virion antibodies in the serum (Fig. 1c). The design of our inactivated vaccine based on this relationship allowed exposure of the N and S antigens, which was determined by visible electron microscopy observation using convalescent serum, mAb-S or mAb-N; the results showed similar interactions of the virion composing the inactivated vaccine and these antibodies (Fig. 1d).

**Figure 1.**
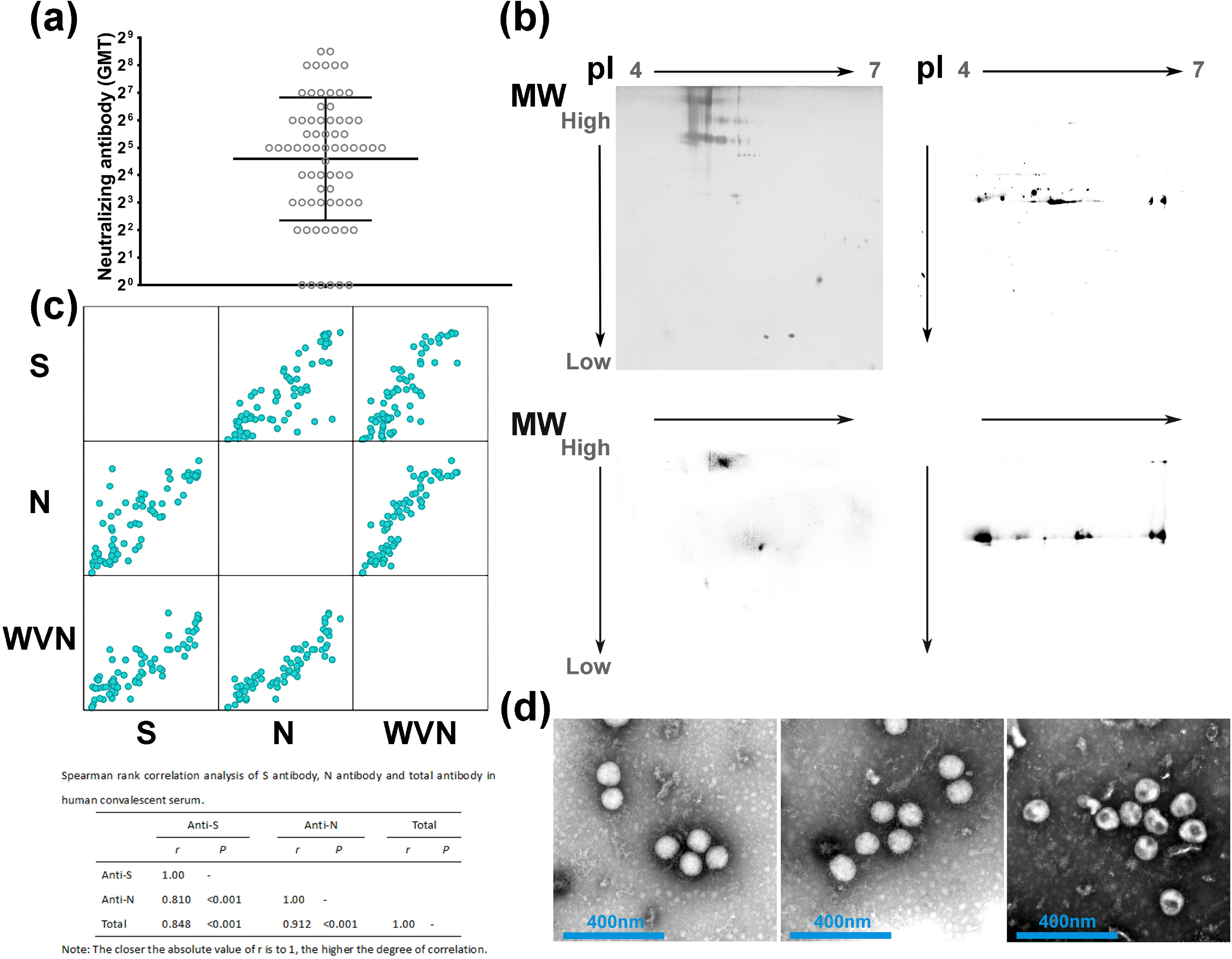
The SARS-CoV-2 inactivated vaccine showed more than one antigenic component recognized by convalescent serum derived from COVID-19 patients. a. The convalescent serum from patients (N=71) could identify and neutralize the virus strain prepared for the SARS-CoV-2 inactivated vaccine. Geometric mean ± SD. b. 2D electrophoresis (upper left) and immune blotting showed the convalescent serum (upper right), mAb-S (lower left) and mAb-N (lower right) could identify the viral proteins in the SARS-CoV-2 inactivated vaccine. c. Correlation analysis of the convalescent serum from patients (N=71) identified the S and N proteins and whole virion (WVN) by Spearman rank correlation analysis. The antibody concentrations were tested and calculated according to a standard curve determined by ELISA. d. The convalescent serum (left), mAb-S (middle) and mAb-N (right) could identify and enrich the virion, as determined by visible electron microscopy.

### Immunization of rhesus macaques with the SARS-CoV-2 inactivated vaccine elicits effective immunity with indexes of humoral and cellular reactions

Based on immunological studies of the SARS-CoV-2 inactivated vaccine in mice, our work basically integrated the GMTs of neutralizing antibodies observed in the convalescent serum from COVID-19 patients and those detected in the serum from mice immunized with the vaccine at various doses (Supplemental Fig. 1). The immune dose of 100 EU was determined to elicit neutralizing antibody titers in the range of 1:16-64 through the intramuscular route with two inoculations on days 0 and 14 (Supplemental Fig. 1). Furthermore, 3 doses of 200, 100 or 20 EU were used to immunize groups A (4 macaques), B (3 macaques) and C (3 macaques), respectively, followed by a booster immunization on the 14^th^ day, while 10 macaques were used as adjuvant controls. The immunological evaluation of these immunized macaques on day 7 after the 2^nd^ inoculation indicated increasing neutralizing antibody titers (Fig. 2a), with GMTs of 107.6, 25.4 and 2 found in the 3 dose groups, respectively, and titers lower than 1:4 were observed in 2 macaques in the low-dose group. This result suggests a dose-dependent relationship for vaccine immunization. However, ELISA analysis with plates coated with the S or N protein indicated no obvious difference in trends of the titers of anti-S and anti-N antibodies (Fig. 2b). Furthermore, ELISPOT analysis of IFN-γ specificity also showed a positive cytotoxic T lymphocyte (CTL) response with no dose difference after stimulation with the S or N antigen (Fig. 2c). These results seem to suggest that the neutralizing antibodies against the S antigen showing a dose-dependent effect related to vaccine immunization are one of the components elicited in the humoral immune reaction.

**Figure 2.**
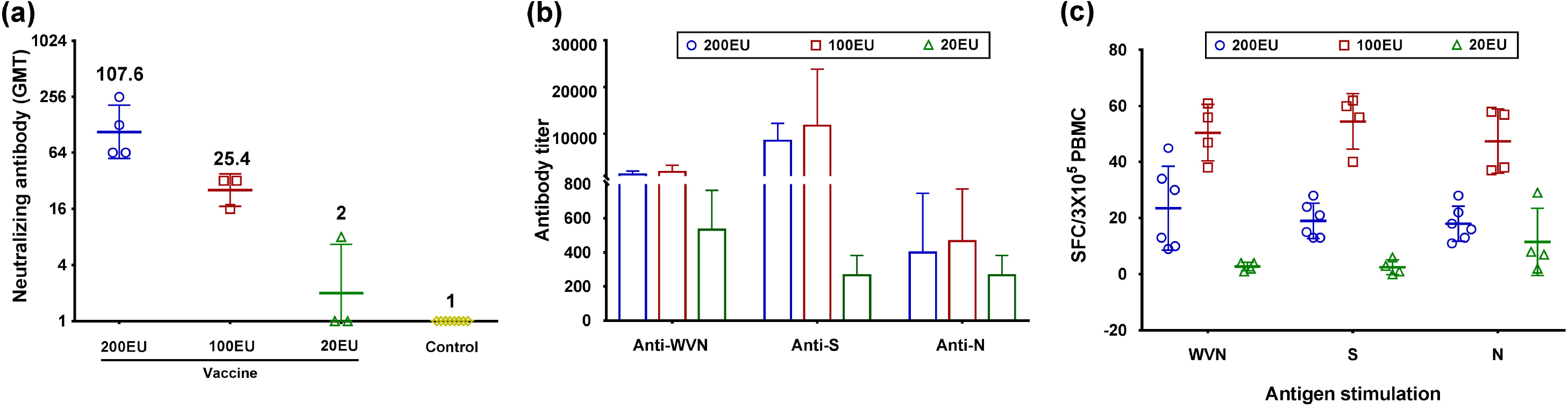
The SARS-CoV-2 inactivated vaccine elicited humoral and CTL immune responses in rhesus monkeys. a. Different doses of the inactivated vaccine induced neutralizing antibodies in rhesus monkeys (N=10). The GMT values for all monkeys in the Al(OH)_3_ adjuvant control group (N=10) were <2 (GMT=1 in the picture). Geometric mean ± SD. b. Different doses of the inactivated vaccine elicited antibodies against the S and N proteins and whole virion (WVN) in rhesus monkeys. The antibody concentrations were tested and calculated according to a standard curve determined by ELISA. Geometric mean ± SD. c. Different doses of the inactivated vaccine elicited IFN-γ-specific immune responses against the S and N proteins and whole virion (WVN) in rhesus monkeys. PBMCs were incubated for 24 h in the presence of a stimulus. Mean ± SD. Samples were obtained on day 7 post 2^nd^ inoculation.

### The integrated immune response elicited by the vaccine is capable of restraining viral replication in the respiratory and alimentary tracts of challenged macaques

Based on the data obtained from above immunological detection of the SARS-CoV-2 inactivated vaccine in rhesus macaques, we designed a viral challenge test involving 10 macaques immunized with the vaccine as described above to identify the immunoprotective effect of the vaccine (Fig. 3a). The other 10 macaques were used as adjuvant controls for observation of clinical manifestations and viral shedding, and one challenged macaque was sacrificed under anesthesia at each time point. Another 2 macaques, which were immunized with a ferritin-fused peptide from the RBD region of the S protein expressed in CHO cells twice with inoculations on days 0 and 14 via the intramuscular route, and their neutralizing antibody titers reached 1:16-32, but no anti-N antibody set-up (Supplemental Fig. 2) was used as a parallel control. All animals were challenged with wild-type virus with dose of 2×10^5^ CCID_50_/each animal via the nasal route. Following viral challenge, monitoring of the body temperature; the viral load in pharyngeal secretions, nasal secretions, and anal swabs; and viremia was performed each day. The results suggested that no obvious fluctuation in body temperature was observed in any immunized macaques, including the two immunized with the peptide vaccine, compared with the 10 adjuvant control animals (Fig. 3b). The detection of viral loads indicated that viral shedding occurred in the nasal cavity of the macaques immunized with the inactivated vaccine, which is the site of virus challenge and the viral load decreased from 36 copies/100 μl - 3 × 10^3^ copies/100 μl on day 1 to less than 50 copies/100 μl on day 2 and was maintained at that level through day 15 post infection (Fig. 3c), while the viral loads in pharyngeal and anal swabs were lower than 50 copies/100 μl (Fig. 3c). Values of 10^3^ - 10^4^ copies/100 μl and higher were found in the pharyngeal, nasal and anal swabs of the adjuvant control macaques for at least 8-9 days (Fig. 3c). In the analysis of the peripheral blood, no positive result was found in the inactivated vaccine group, but a peak value was observed on days 5-7 in the positive control group (Fig. 3c). Interestingly, the two macaques immunized with the peptide vaccine had neutralizing antibody titers of 1:16-32 and showed values higher than 10^4^ and 10^3^ copies/100 μl in nasal and anal swabs, respectively, on days 3-5 and 6-8 (Fig. 3c). These results suggest not only that the inactivated vaccine elicits a valid immunoprotective effect but also that an integrated immune response rather than neutralizing antibodies alone may be needed to restrain viral replication in the respiratory and/or alimentary tracts.

**Figure 3.**
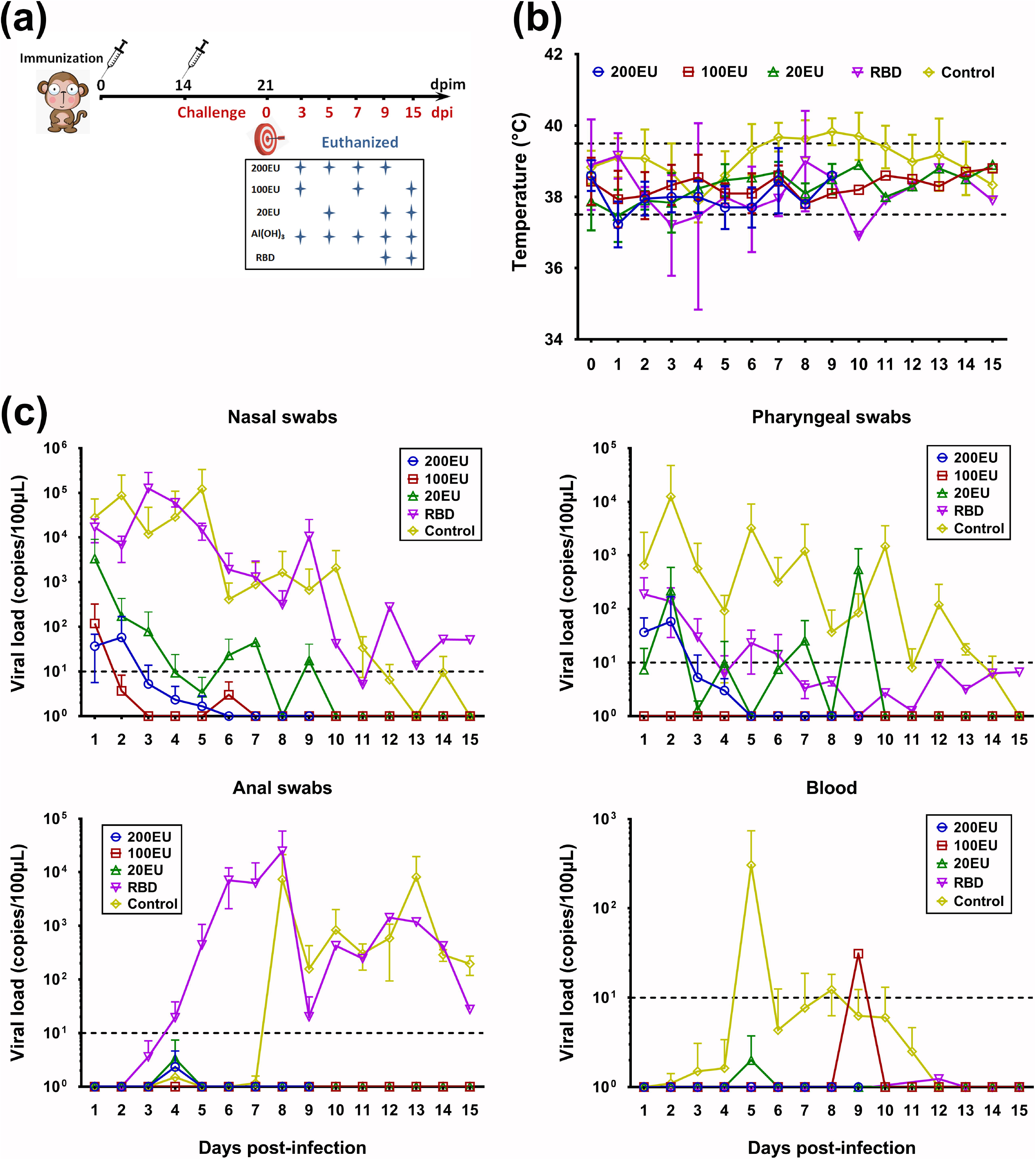
The integrated immune response elicited by the inactivated vaccine can restrain viral replication in the respiratory and/or alimentary tracts. a. The design of the immunity and viral challenge protocol. b. Monitoring of the body temperature of rhesus monkeys infected by SARS-CoV-2. The normal range from 37.5 to 39.5°C (refer to the normal monkeys (N=10) monitored during the same period) is indicated with dotted lines. Inactivated vaccine (200EU, 100EU and 20EU; N=10), RBD peptide vaccine (N=2) or Al(OH)_3_ adjuvant (control; N=10). Mean ± SD. c. The viral loads of pharyngeal secretions, nasal secretions, anal swabs and blood from monkeys immunized with the inactivated vaccine (200EU, 100EU and 20EU; N=10), RBD peptide vaccine (N=2) or Al(OH)_3_ adjuvant (control; N=10) after infection with live virus. Negative is the copies less than 10 (dotted lines). Mean ± SD.

### The immunity induced by the inactivated vaccine can disrupt viral proliferation in organs and largely alleviate pathological damage to tissues

Generally, the experimental index of immunoprotective effects on animal models for the study of inactivated virus vaccines includes viral replication and the virus-induced damage to tissues of vaccine-immunized animals during challenge tests with a wild-type virus (21, 22). Our work here also focused on the detection of the viral loads in tissues and pathological lesions in these tissues in immunized macaques, which were sacrificed under anesthesia on day 3, 5, 7, 9, or 15 post viral challenge. All tissue samples collected from these challenged animals, including adjuvant control monkeys and the monkeys immunized with the RBD peptide vaccine, were used for the detection of viral loads and pathological observation. q-RT-PCR results suggested that almost all samples from various tissues of the animals in the inactivated vaccine immunization group showed values lower than 50 copies/100 mg and only those in spleens were higher than 100 copies/100 mg (Fig. 4a). Macaque No. 19177, which was immunized with the RBD peptide vaccine and sacrificed under anesthesia on day 9, was found to have 635, 2200 and 140 copies/100 mg in the intestine, cervical lymph nodes and spleen, respectively (Fig. 4a); and macaque No. 19295, which was in the same group and sacrificed under anesthesia on day 15, was found to have 288,670 copies/100 mg in the cerebellum (Fig. 4a). However, the evaluation of adjuvant control animals showed a trend toward increasing viral loads in various organs during the observation period (Fig. 4a). In the pathological detection at 5 time points, the lung tissues of 3 groups of macaques immunized with the inactivated vaccine showed slight and nonspecific inflammatory reactions, including a few local aggregations of inflammatory cells, inflammatory exudation in some alveoli and bronchioles, slight hyperplasia of the epithelial tissue in a few alveolar tissues and slight congestive reactions (Fig. 4b). The two macaques immunized with the RBD peptide vaccine presented similar changes (Fig. 4b), while the macaques in the adjuvant control group presented more severe inflammatory reactions, which were recognized as interstitial pneumonia based upon the histopathologic observation (Fig. 4b). These results suggest that the immunity elicited by the inactivated vaccine can restrain viral proliferation in various tissues in immunized individuals and alleviate pathological lesions in tissues caused by viral replication. Interestingly, elimination of the challenge virus by this immunity in macaques did not seem to be due to the existence of neutralizing antibodies. Two macaques immunized with the inactivated vaccine with a neutralizing antibody titer lower than 1:4 were capable of completely restraining viral replication *in vivo*, but the other two immunized with the RBD peptide vaccine had a titer of 1:16-32 and were not capable of completely eliminating viral replication.

**Figure 4.**
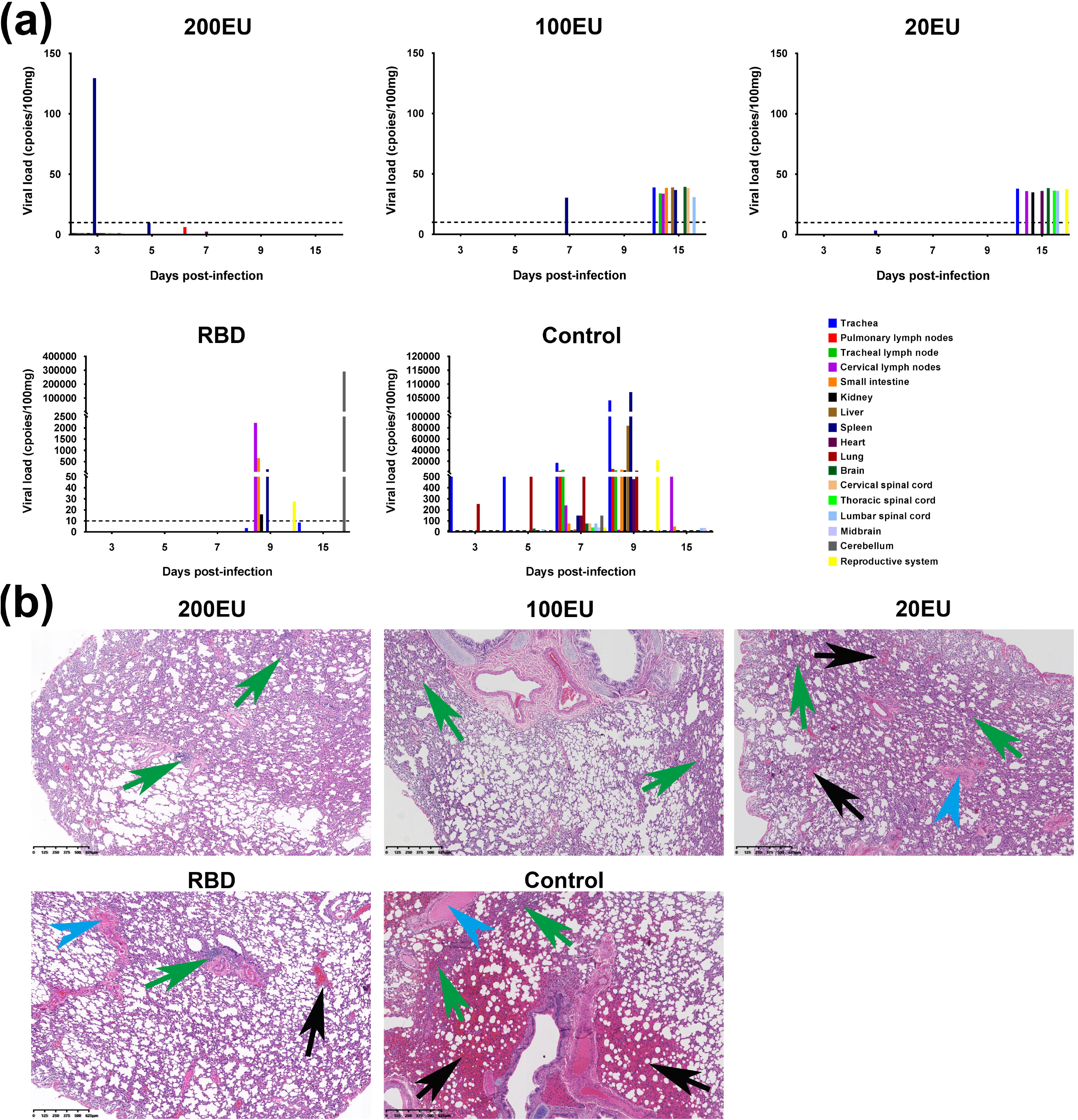
The integrated immune response elicited by the inactivated vaccine can eliminate viruses and alleviate the pathological damage induced by the challenge virus. a. The viral loads of various organs from monkeys immunized with the inactivated vaccine (200EU, 100EU and 20EU; N=10), RBD peptide vaccine (N=2) or Al(OH)_3_ adjuvant (control; N=5) after infection with live virus. Negative is the copies less than 10 (dotted lines). Mean ± SD. There are no data on day 15 of the 200EU group, on days 5 and 9 of the 100EU group, on days 3 and 7 of the 20EU group, and on days 3, 5, and 7 of the RBD group. b. Typical pathological changes in the lungs of rhesus monkeys immunized with the inactivated vaccine (200EU, 100EU and 20EU; N=10), RBD peptide vaccine (N=2) or Al(OH)_3_ adjuvant (control; N=5) and infected with live virus. Congestion (black arrow), infiltration of inflammatory cells (green arrow) and edema (blue arrow). Samples were obtained at 7 dpi (200EU), 7 dpi (100EU), 9 dpi (20EU), 9 dpi (RBD vaccine) and 7 dpi (control).

### The immune response elicited in macaques by the inactivated vaccine presents an immunologically dynamic process

Previous reports on COVID-19 suggest it to be an acute and severe inflammatory disease of the respiratory system (1, 23) and lead to the deduction that the pathological mechanism may be due to not only viral infection but also a process of excessive immune reaction, especially ADE, induced by the virus (12). In this case, the study of the SARS-CoV-2 inactivated vaccine in an animal model should be focused on the immunologically dynamic process associated with the elicitation of specific antiviral immunity through monitoring various indexes of the immune cell population, which should be maintained in a stable dynamic state (24). Our work detected altered proportions of different immune cells in the total population of PBMCs after viral challenge in macaques immunized with the inactivated vaccine. The results suggested that compared to that of the adjuvant control animals (Fig. 5a), the dynamic alteration in the PBMC population in all animals immunized with inactivated vaccine included increases in the proportions of T cells, NK cells, T cells with IFN-γ specificity and Treg cells (Fig. 5a, b), in which upregulation of the percentages of IFN-γ-specific T cells and Treg cells was observed on days 9 and 5 post viral challenge, respectively (Fig. 5b). Furthermore, the detection of some inflammatory factors, including IL-2, IL-4, IL-5, IL-6 and TNF-α, in the serum of all animals indicated that viral challenge was capable of slightly upregulating IL-2 and IL-5 levels in the inactivated vaccine group (Fig. 5c). These results suggest not only the dynamic activation of the immune system induced in macaques immunized with the inactivated vaccine.

**Figure 5.**
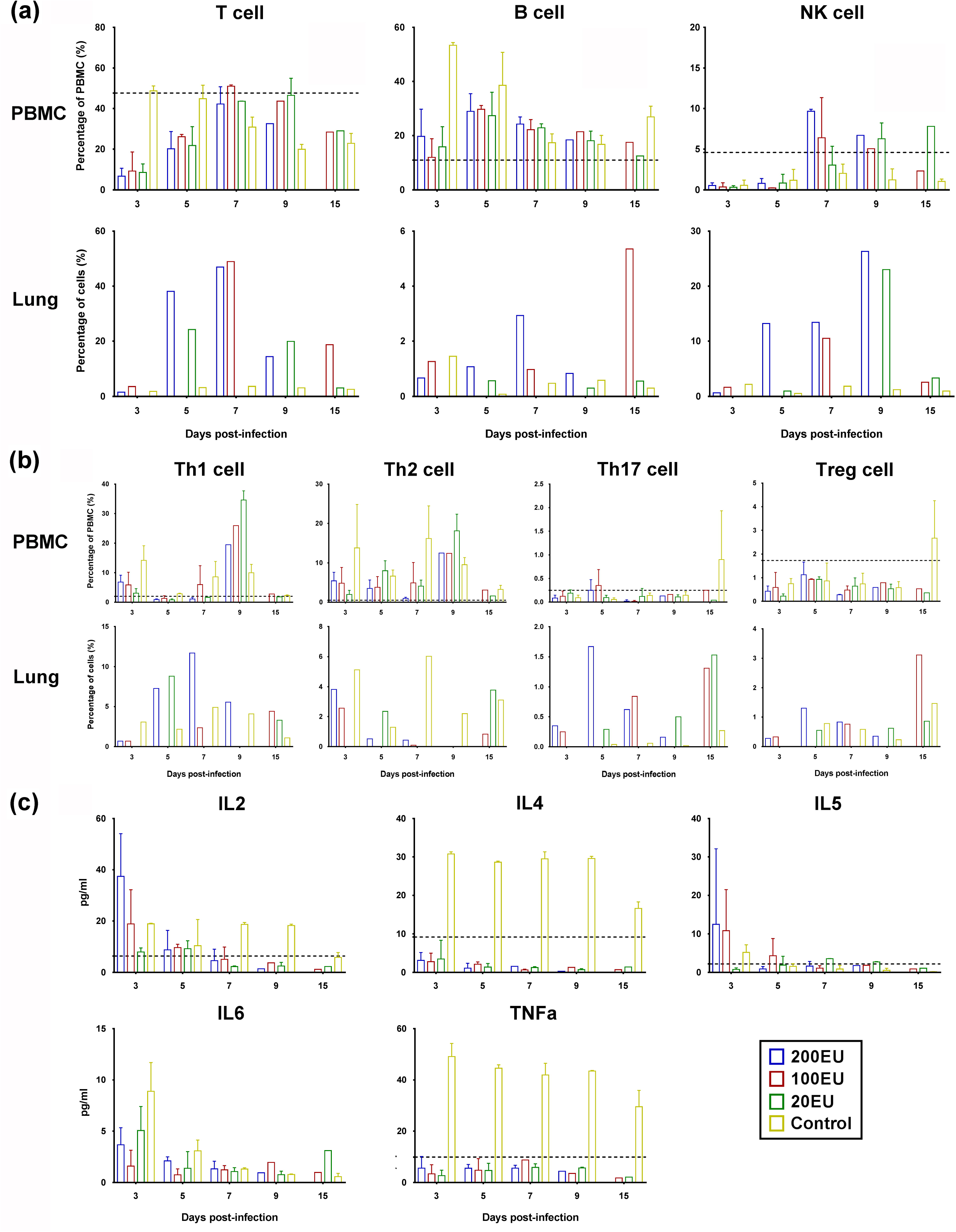
The dynamic immune reaction process is maintained in macaques immunized with the inactivated vaccine during viral challenge. a. The percentages of T, NK and B cells in PBMCs or lung from rhesus monkeys immunized with the inactivated vaccine (N=10) or Al(OH)_3_ adjuvant (control; N=5) during viral challenge. Mean ± SD. b. The percentages of Th1, Th2, Th17 and Treg cells in PBMCs or lung from rhesus monkeys immunized with the inactivated vaccine (N=10) or Al(OH)_3_ adjuvant (control; N=5) during viral challenge. Mean ± SD. c. IL-2, IL-4, IL-5, IL-6 and TNF-α levels in the serum of rhesus monkeys immunized with the inactivated vaccine (N=10) or Al(OH)_3_ adjuvant (control; N=5) during viral challenge. Mean ± SD. The normal range (refer to the normal monkeys (N=10) monitored during the same period) is indicated with dotted lines.

## Discussion

The global public health crisis induced by SARS-CoV-2, which leads to acute and severe infectious respiratory disease and/or pneumonia, is driving the rapid development of a suitable vaccine (3, 25). Facing this emergency requirement, a basic scientific question that first needs to be addressed is the antigen composition capable of eliciting effective immunity in a vaccine; however, no clear antigenic analysis of different viral structural proteins is available. We investigated the interaction of our inactivated vaccine and convalescent serum from COVID-19 patients based on the deduction that the recovery of a patient means his/her immune system was capable of controlling and/or eliminating the virus, and convalescent serum may contain antibodies effective in this process. The data from this study suggest that anti-S and anti-N antibodies may play similar roles in convalescent serum and that both types of antibodies, but especially anti-N antibodies, show a strong relationship with the general antibody response to the virus. This result led to our technical design of an inactivated vaccine capable of exposing the S protein and N protein upon immunization into animals. In the viral challenge test with macaques, this vaccine elicited a specific immune response with a neutralizing antibody titer that increased depending on the dose and specific CTL response against the S-antigen, N-antigen and virion. The ELISA confirmed that both anti-S and anti-N antibodies were elicited in these animals, which is different from the serum of animals immunized with the RBD peptide vaccine containing only anti-S antibodies that showed neutralizing activity of 1:16-32. Usually, neutralizing antibodies are capable of blocking the binding of a virus to receptors and are an index indicative of effective antiviral immunity (26, 27). In our study of the inactivated vaccine, an immunoprotective effect against viral challenge was observed in macaques immunized with high and medium doses of the inactivated vaccine, which were associated with increasing neutralizing antibody titers, while similar protective effects were observed in low-dose immunized animals with a neutralizing antibody GMT of 2 and a CTL response similar to those of the high- and medium-dose groups. Interestingly, the immunity elicited in macaques immunized with the RBD peptide vaccine was unable to completely restrain viral proliferation in some tissues, even though the animals possessed neutralizing antibodies with titer of 1:16-32. Two animals were found to have high viral shedding in the nasopharynx and/or alimentary tract during later times of viral infection with slightly more serious pathological inflammatory reactions in the lungs than animals in the high- and medium-dose groups treated with inactivated vaccine immunization. These results suggest that the immunity elicited by our inactivated vaccine can provide systematic immune protection against viral infection not only by producing upregulation of neutralizing antibody titers but also by functioning through the CTL response associated with increased anti-N antibody levels. This conclusion that integrated immunity involving antibodies against various viral proteins elicited by an inactivated vaccine is needed for the effective prevention of SARS-CoV-2 infection, at least in macaques, is supported by observation of macaques immunized with the RBD peptide vaccine. All data obtained here may lead to a logical analysis of the interaction of SARS-CoV-2 with the immune system, which indicates that the N protein may play an important role in the activation of innate immunity in epithelial cells to further the specific antiviral immune response via the N protein being a major viral antigen that interacts with PRRs in cells, while the S protein greatly impacts the elicitation of neutralizing antibodies and interacts with PRRs to a lesser degree than the N protein. In this case, a valid systematic immunoprotective response against SARS-CoV-2 infection should include at least anti-S and anti-N antibodies, and innate immunity may contribute largely to this response and should be considered in vaccine development. The technical design of our inactivated vaccine enables N and S antigen exposure and elicits antibodies against both viral proteins, which indicates that this vaccine is of practicable significance. However, the observation of immunized macaques with a low titer of neutralizing antibodies due to the very low antigen amount does not suggest that ADE exists. Based on these data, this vaccine was permitted to enter clinical trial by CFDA with approval number of 2020L00020.

## Methods

### Virus and cells

The SARS-CoV-2 virus used was isolated from the respiratory secretions of an adult male patient at Yunnan Hospital of Infectious Diseases in Kunming in January 2020. The virus proliferated in Vero cells (ATCC, Manassas, USA) and was purified by plaque cloning. The cloned virus was identified via genomic sequencing and named KMS-1 (GenBank No: MT226610.1). Vero cells were cultured in DMEM (Corning, NY, USA) containing 5% fetal bovine serum (FCS; HyClone, Logan, USA).

### Viral titration

All experimental procedures were performed under BSL-3 laboratory conditions. Virus samples were serially diluted 10-fold with serum-free DMEM (Corning, NY, USA). Different dilutions of the virus were added to a 96-well plate. Each dilution (100 μl per well) was added to 8 parallel wells. Then, 100 μl of Vero cell suspension was added to each well at a concentration of 2.5 × 10^5^ cells/ml. After the plate was incubated at 37°C in 5% CO_2_ for 7 days, the cytopathic effect (CPE) was observed and assessed with an inverted microscope (Nikon, Tokyo, Japan).

### Inactivated vaccine

The SARS-CoV-2 inactivated vaccine was developed by the Institute of Medical Biology (IMB), Chinese Academy of Medical Sciences (CAMS). Briefly, the virus seed strain for the vaccine was inoculated into Vero cells, which was provided by WHO with batch number of UCC91-02 of main seed for vaccine production, and obtained the verification approval number of 201501 from National Institute for Food and Drug Control (NIFDC) of China, in BSL-3 environment. The viral harvest was inactivated by formaldehyde in rate of 1:4000 for 48 hours, which was found enable to break viral membrane. After this inactivation, the chromatograph process using Core-700 gel medium was performed for the first purification. Further, viral antigenic component collected was concentrated for the second inactivated with beta-propiolactone in rate of 1:2000 destructing viral genomic structure. After the second purification by the same medium, the vaccine stock was evaluated with various quality indexes including antigen content, immunogenicity, sterility and residues test etc. The viral antigen content was measured via ELISA assay, in which, the antibodies against S, N protein and whole virion were used to coat plate for the detections of each antigenic component. A solution of the inactivated virus vaccine stock was then emulsified in 0.5 mg/ml Al(OH)_3_ adjuvant, constituting the final vaccine.

### Ethics

#### Human

Convalescent serum samples were collected from patients diagnosed with new coronavirus pneumonia at Yunnan Hospital of Infectious Diseases, Kunming Third People’s Hospital and CDC of Xianyang city, Hubei province, with the patients providing informed consent. The protocols were reviewed and approved by the Experimental Management Association of the IMB, CAMS (approval number: DWSP 202003 004).

#### Animal

The animal experiment was designed and performed according to the principles in the ‘‘Guide for the Care and Use of Laboratory Animals” and in ‘‘Guidance for Experimental Animal Welfare and Ethical Treatment”. The protocols were reviewed and approved by the Experimental Animal Management Association of the IMB, CAMS (approval number: DWSP 202003 005). All animals were fully under the care of veterinarians at the IMB, CAMS.

### Animals

#### Mouse

Four-week-old female BALB/c mice (Beijing Vital River Laboratory Animal Technologies Co. Ltd, Beijing, China) were housed in a laboratory (ABSL-3) within the specific pathogen-free facility at the IMB, CAMS. Mice were anesthetized with inhaled 2% isoflurane for all procedures, with every effort made to minimize suffering.

#### Monkey

Rhesus monkeys (age 1.5-2 years) were bred and fed pellets (IMB, CAMS, China) and fresh fruits in a laboratory (ABSL-3) at the IMB, CAMS.

### Immunization

#### Mouse

Mice were randomly divided into four groups, intramuscularly immunized with the vaccine at 200 ELISA units (EU; viral antigen concentration determined by ELISA), 100 EU or 20 EU or with Al(OH)_3_ adjuvant alone, which was used as a control, on days 0 and 14. Blood samples were collected for neutralization assays on day 7 after the booster injection.

#### Monkey

Rhesus monkeys were randomly divided into four groups intramuscularly immunized with the vaccine at 200 EU (N=4), 100 EU (N=3) or 20 EU (N=3) or with Al(OH)_3_ adjuvant alone (N=10) as a control on days 0 and 14. The monkeys were tested for neutralizing antibodies and IFN-γ-secreting cells in blood samples taken on day 7 after the booster injection.

### Viral challenge

SARS-CoV-2 infection (2×10^5^ CCID_50_/monkey) via nasal spray was performed under ABSL-3 laboratory conditions for the immunized macaques by inactivated vaccine and adjuvant control group. As a parallel control, two monkeys immunized with a peptide vaccine (Supplemental Fig. 2) were also infected with the patient-derived live SARS-CoV-2 strain. All animals were monitored daily for clinical signs. Pharyngeal secretion, nasal secretion, anal swab and blood samples were obtained every day after infection. Blood was collected under appropriate anesthesia to alleviate pain and minimize suffering.

In the 200 EU-dose vaccine group, a monkey was euthanized on days 3, 5, 7 and 9 postinfection (dpi). In the 100 EU-dose vaccine group, a monkey was euthanized on days 3, 7 and 15 dpi. In the 20 EU-dose vaccine group, a monkey was euthanized on days 5, 9 and 15 dpi. In the adjuvant control group, a monkey was euthanized at each time point (days 3, 5, 7, 9 and 15 dpi). In the RBD peptide vaccine group, a monkey was euthanized on days 9 and 15 dpi. The blood and organs of the euthanized monkeys were obtained for various experiments. The remaining 5 macaques in the adjuvant control groups were used for observation of clinical manifestations and viral shedding.

### 2D protein electrophoresis-Western blot analysis

Purified virus samples were resuspended in 2D lysis buffer (8 M urea, 2 M thiourea, 4% CHAPS, 100 mM DTT, and 2% IPG buffer). Total protein was quantified using the PlusOne 2D Quant Kit (GE Healthcare Europe GmbH, Freiburg, Germany) according to the manufacturer’s instructions. Approximately 200 μg of protein were first separated based on their pI using immobilized linear gradient strips (ImmobilineTM DryStrip, Amersham Biosciences Europe GmbH, Freiburg, Germany) covering the pH range 4-7 and then separated by 12% SDS-PAGE. Proteins were transferred to a polyvinylidene fluoride membrane. The transferred membrane was blocked in 5% bovine serum albumin (BSA)-Tris-buffered saline/Tween 20 (Tris-HCl, 100 mM, pH 7.5; NaCl, 0.9%; and Tween-20, 0.2%) and treated with convalescent serum, anti-S (Snio Biological, Beijing, China), anti-N antibody (Snio Biological) and horseradish peroxidase (HRP)-labeled secondary antibody (Abcam, MA, USA) to visualize the proteins according to the standard protocol of the enhanced chemiluminescence (ECL) reagent.

### Neutralizing antibody test

Heat-inactivated serum was diluted and coincubated with live virus (100 lgCCID_50_/well) for 2 h at 37°C, followed by addition of Vero cells (10^5^/mL) of 100 μl to the mixture. Then, the plates were incubated at 37°C in 5% CO2 for 7 days. The CPEs were observed and assessed with an inverted microscope (Nikon) to determine the neutralizing antibody titer of the serum. The geometric mean titers (GMTs) of neutralizing antibodies were measured.

### ELISA

The S protein (Sanyou Biopharmaceuticals Co., Ltd., Shanghai, China), the N protein (Sanyou Biopharmaceuticals Co., Ltd.) and purified viral antigen were used to coat separate wells of 96-well ELISA plates (Corning, NY, USA) at a concentration of 5 μg/well and incubated at 4°C overnight. Then, the plates were blocked with 5% BSA-phosphate-buffered saline (PBS), incubated with serum samples, and visualized with an HRP-conjugated antibody (Abcam, MA, USA) and TMB substrate (Solarbio, Beijing, China) according to previously described methods (28). The absorbance of each well at 450 nm was measured using an ELISA plate reader (Gene Company, Beijing, China), and the following equation was used: resulting OD = (experimental well OD) – (mock well OD). The standards used were commercial antibodies at known concentrations (Snio Biological). Antibody concentrations were calculated according to a standard curve.

### ELISPOT

An ELISPOT assay was performed with the Monkey IFN-γ ELISPOT Kit (Mabtech, Cincinnati, OH, USA) according to the manufacturer’s protocol. Briefly, peripheral blood mononuclear cells (PBMCs) were isolated from the blood by lymphocyte isolation (Ficoll-Paque PREMIUM; GE Healthcare, Piscataway, NJ, USA) and plated in duplicate wells. Three different stimulators, purified SARS-CoV-2 antigen, recombinant S-protein (Sanyou Biopharmaceuticals Co., Ltd.) and recombinant N-protein (Sanyou Biopharmaceuticals Co., Ltd.), were added to separate wells. The positive control was phytohemagglutinin (PHA). The plate was incubated at 37°C for 24 h, after which time the cells were removed, and the spots were developed. The colored spots were counted with an ELISPOT reader (CTL, Shaker Heights, OH, USA).

### Electron microscopy

Purified inactivated SARS-CoV-2 preparations were coincubated with convalescent serum, a monoclonal antibody (mAb) against S (mAb-S) or N (mAb-N) (Solarbio, Beijing, China) at 37°C for 24 h, stained with 1% phosphotungstic acid and observed by transmission electron microscopy (Hitachi, Kyoto, Japan).

### Quantitation of the viral load by q-RT-PCR

Total RNA was extracted from blood and tissue samples from experimental monkeys with TRIzol reagent (Tiangeng, Beijing, China). According to the protocol, q-RT-PCR was performed using the One Step PrimeScript™ RT-PCR Kit (Perfect Real Time; TaKaRa). The primers used for q-RT-PCR were selected to be specific for the E and ORF1ab sequences in the SARS-CoV-2 genome (Table 1). Viral copies were quantified according to vitro-synthesized RNA, and the quantity was expressed as a relative copy number, determined by the equation [(μg of RNA/μl) / (molecular weight)] × Avogadro’s number = viral copy number/μl (29).

**Table 1.**
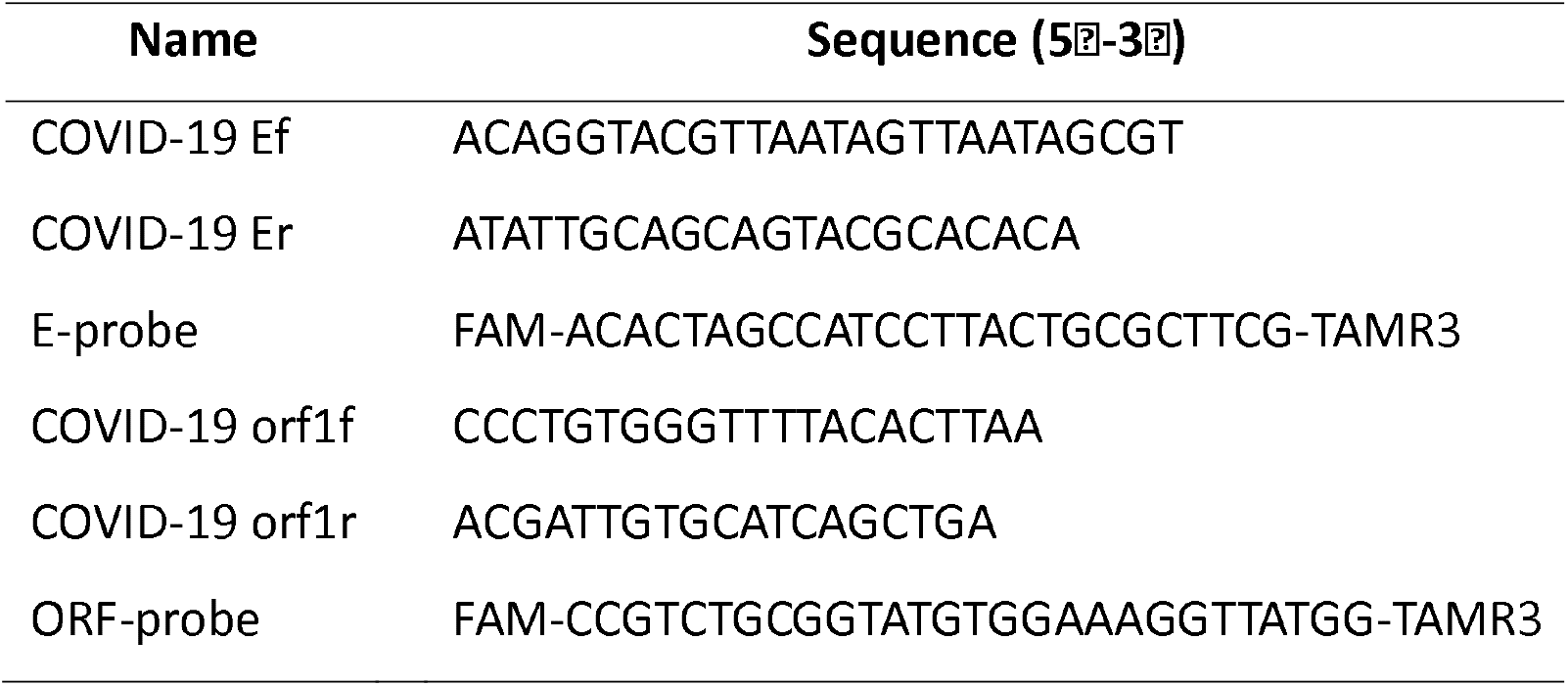
The primers of qRT-PCR to detect SARS-CoV-2.

### Histopathological examinations

The organs of experimental monkeys were fixed in 10% formalin, embedded in paraffin, sliced into 4-μm sections and stained with hematoxylin and eosin (H&E). Morphology was assessed with an inverted microscope (Nikon).

### Immune cell populations

PBMCs were isolated from monkeys by lymphocyte isolation (Ficoll-Paque PREMIUM; GE Healthcare). Anti-CD3, anti-CD20 and anti-CD16 antibodies were added to the PBMCs. The mixtures were incubated at room temperature (RT) for 30 min in the dark. Reagents for red blood cell lysis (BD) and membrane permeabilization (BD) were added in sequence. After washing with PBS twice, the cells were resuspended in PBS and detected using a flow cytometer (BD, USA). T cells (CD3^+^), B cells (CD20^+^) and NK cells (CD16^+^) were evaluated. Furthermore, the T cells were typed into T helper (Th) 1, Th2, regulatory T (Treg) and Th17 cells. The PBMCs were coincubated with an anti-CD4 antibody (BD) at RT for 30 min. Red blood cell lysis (BD) and membrane permeabilization (BD) reagents were added. After incubating with the red blood cell lysis and membrane permeabilization reagents and washing with PBS twice, the cells were stained with anti-FOXP3, anti-IL-4, anti-IFN-γ and anti-IL-17A antibodies for 30 min at RT and then washed once. The percentages of immune cells were detected using a flow cytometer (BD). Th1 cells (CD4^+^/IFN-γ^+^), Th2 cells (CD4^+^/IL-4^+^), Treg cells (CD4^+^/FOXP3^+^) and Th17 cells (CD4^+^/IL-17A^+^) were assessed.

### Cytokine assay

The levels of IL-2, IL-4, IL-5, IL-6, TNF-α and IFN-γ in the serum of experimental monkeys were detected with the Non-Human Primate (NHP) Th1/Th2 Cytokine Kit (BD) according to the manufacturer’s protocol. Briefly, serum was added into a tube containing detection beads. Then, the PE detection reagent was added, and the mixtures were incubated at RT for 2 h in the dark. After washing, the beads were resuspended in wash buffer and detected using a flow cytometer (BD). The levels of these cytokines were calculated according to a standard curve.

### Statistical analysis

Data are shown as the mean and standard deviation. GraphPad Prism software (San Diego, CA, USA) and STATA (Version 15.0; STATA Corp., College Station, TX, USA) were used for statistical analyses. The association of antibodies against S, N and virus (WVN) was Spearman rank correlation analysis.

## Author contributions

QHL, LDL and CGL conceived and designed the experiments; HBC, ZPX, RXL, STF, HL, ZLH, KWX, YLiao, LCW, XQL, TWM, YYY, LG, JBY, HWZ, XLX, JL, YLiang, DDL, HZ, GRJ, FMY, YGH, XW, CC, XQD, YLi, HLZ, JZ, HJY, JFY and WLZ performed the experiments; QHL, LDL, YZ and ZMZ analyzed the data; XQD, XFZ, CH, YCC, JP, KLM, LW, WGD, DS, HLZ, RJJ, LY, MJX, LYi, ZXZ, ND, HYang and WY contributed reagents/materials/analysis tools; and QHL wrote the paper.

## Acknowledgments

This work was supported by grant of National Key R&D Program of China (2020YFC0849700) and Major science and technology special projects of Yunnan Province.

