## Supplemental Figure 1 for "A valid protective immune response elicited in rhesus macaques by an inactivated vaccine is capable of defending against SARS-CoV-2 infection"

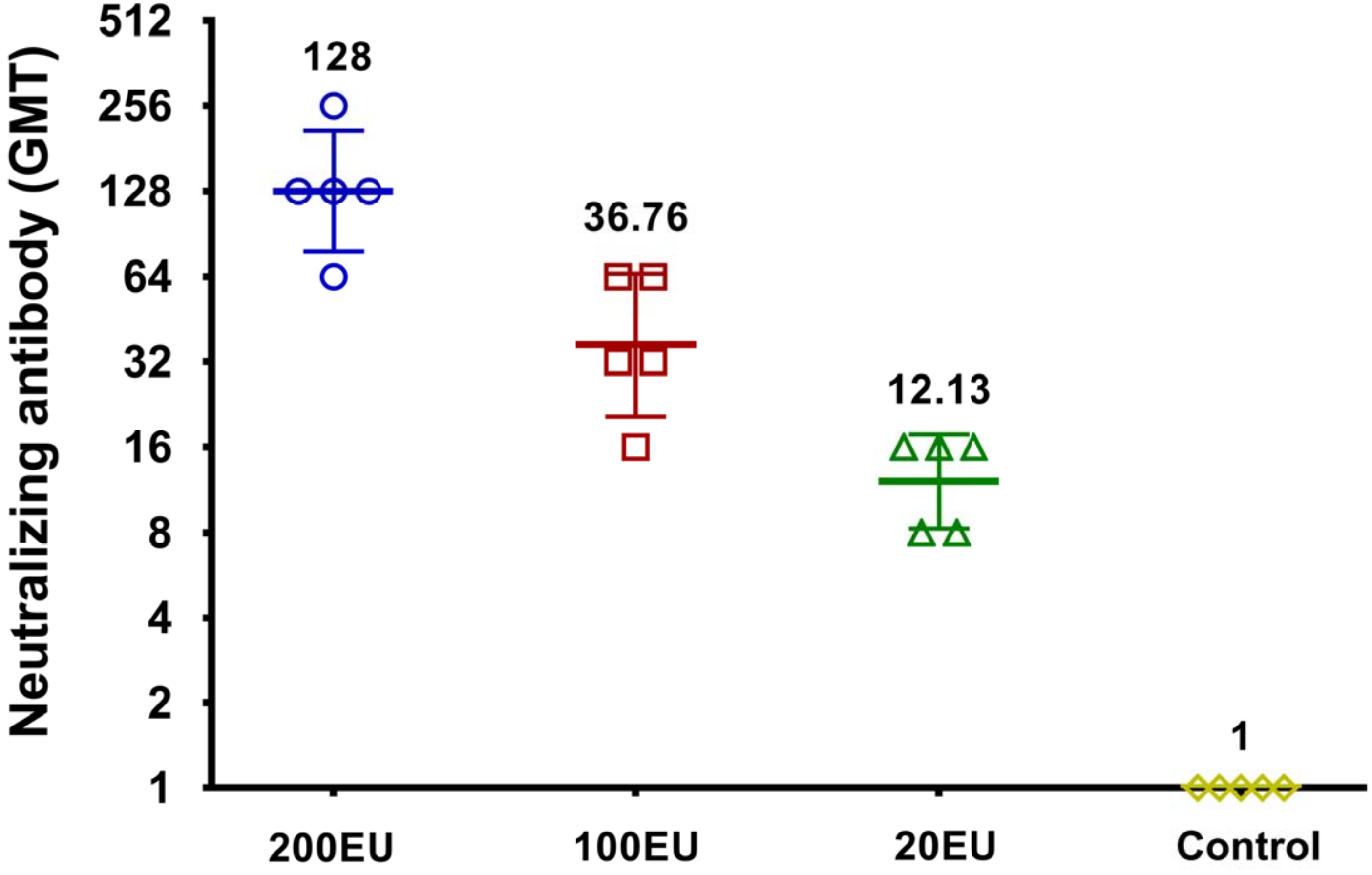


**Figure S1. Different doses of the SARS-CoV-2 inactivated vaccine induced neutralizing antibodies in mice.**

The GMT values for the control ($Al(OH)_3$ adjuvant) groups were all < 2. 200EU (N=10), 100EU (N=10) and 20EU (N=10). Geometric mean ± SD.
