## Supplementary figures and images for "A valid protective immune response elicited in rhesus macaques by an inactivated vaccine is capable of defending against SARS-CoV-2 infection"

### Supplemental Figure 2

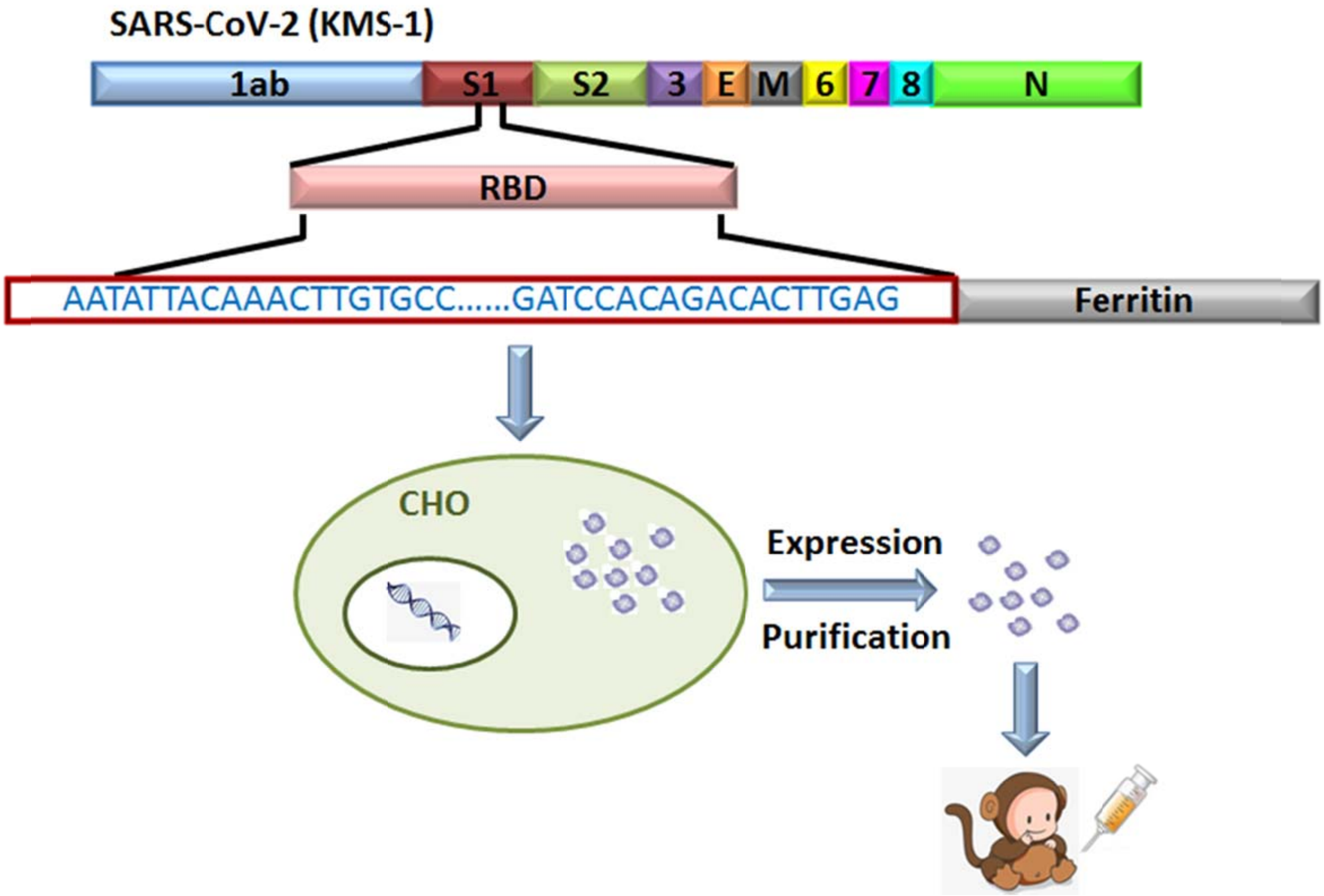


**Figure S2. Development of the RBD peptide vaccine**
