## Supplemental Figure 3 for "A valid protective immune response elicited in rhesus macaques by an inactivated vaccine is capable of defending against SARS-CoV-2 infection"

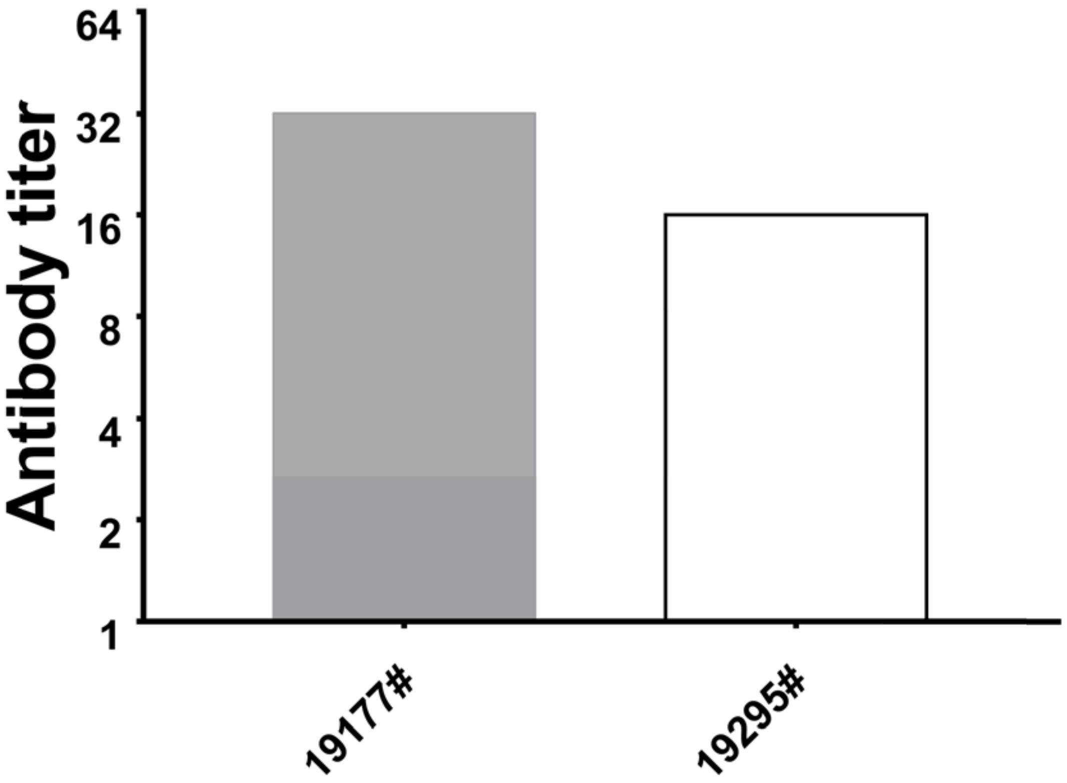


**Figure S3. The SARS-CoV-2 RBD peptide vaccine elicited humoral immune responses in rhesus monkeys**

Neutralizing antibodies were elicited in rhesus monkeys.
